# Multivariate pattern classification on BOLD activation pattern induced by deep brain stimulation in motor, associative, and limbic brain networks

**DOI:** 10.1101/574889

**Authors:** Shinho Cho, Hoon-Ki Min, William S. Gibson, Myung-Ho In, Kendall H. Lee, Hang Joon Jo

## Abstract

Functional magnetic resonance imaging (fMRI) concurrently conducted with the deep brain stimulation (DBS) has shown that diffuse BOLD activation occurred not only near stimulation locus, but in multiple brain networks, supporting that network-wide modulation would underlie its therapeutic effect. While the extent and pattern of activation varies depending on specific anatomical locus stimulated by DBS, some stimulation targets could induce similar activation pattern in cerebral cortex, albeit different therapeutic and adverse effects were yielded.

In order to characterize the unique network-level activation effects of three DBS targets (subthalamic nucleus, the globus pallidus internus, and the nucleus accumbens), we trained the pattern classifier with DBS-fMRI data from three stimulation groups (21 healthy swine), wherein five six seconds of electrical stimulation was conducted while gradient-echo echo planar imaging was on going. Then whole brain regions were systematically grouped into different size of network-of-interest and the classification accuracy for individual target region was quantitatively assessed. We demonstrated that the pattern classifier could successfully differentiate BOLD activation pattern of cortical and subcortical brain regions originated from each individual stimulation target. Moreover, the success rate of classification indicated that some brain regions evoked indistinguishable BOLD pattern, suggesting the presence of commonly activated regions, which was influenced by stimulating different DBS targets.

Our results provide an understanding of the biomarker of BOLD pattern that is associated with clinical effectiveness as well as an adverse effect associated to the stimulation. Further, we provide the proof-of-concept for multivariate pattern analysis that is capable of disentangling the complicated BOLD activation pattern, which cannot be readily achieved by a conventional univariate analysis.

## INTRODUCTION

Stimulating human subcortical structures via the Deep brain stimulation (DBS) have been showing clinical efficacy [1–3]. For example, the subthalamic nucleus (STN) and globus pallidus internus (GPi) have been popular targets for DBS surgery, since those has been known for alleviating motor-related symptoms in Parkinson’s disease and essential tremor. Furthermore, recent DBS application is being extended to treat or alleviate neuropsychological disorders, for example, obsessive-compulsive disorder (ODC) with stimulating nucleus accumbens (NAc) targets. [4–6].

Up to date, the activation mechanism of DBS has been investigated at the cellular and local circuitry levels, but it has been poorly understood that how selecting a specific DBS target modulates distinct distal and diffuse functional networks. While stimulation was initially believed to inhibit the pathological transmission in neural circuits near the stimulation locus (‘jamming’) [7, 8], recent evidence from electrophysiological [4, 9], modeling [10] and functional imaging studies [11, 12] suggest that DBS effect from any given stimulation locus, in fact, diffuses across a wide array of limbic, cognitive, and motor-related functional networks (‘network effect’) [13–15].

In particular, recent functional magnetic resonance imaging (fMRI) during DBS has demonstrated that DBS evokes characteristic, but spatially overlapped blood-oxygenated-level-dependent (BOLD) activation pattern across a brain. For example, STN and GPi stimulation both induces BOLD response in the premotor cortex, striatum, and thalamus in humans and swine [16–18]. STN and NAc DBS yields non-motor, reward-related brain regions in human subjects [19] and in a swine model [20].

Despite many attempts of DBS-fMRI studies to reveal the unique activation pattern associated to different DBS targets, it has been yet clear which network-level neuromodulatory mechanism underlies on therapeutic and adverse effect. Although many possible reasons may exist, it must be noted that spatially overlapped BOLD patterns could be found between different DBS targets [21]. Furthermore, some of these distinctive targets have been showing similar therapeutic effect, but also believed to be associated with different adverse effects. Up to date, data analysis of fMRI mostly adopted putatively called “massively univariate” approach [22] of region-of-interests (ROIs) or population of voxels. However, more recently multivariate approach, i.e., multivariate pattern classification, have suggested that conventional analyses could be insensitive to detect distinctiveness between spatial patterns of BOLD activation. Thus, which extent the similarity and dissimilarity of DBS-evoked modulation pattern effects have not been fully revealed for a given DBS targets, which mediates behavioral and clinical expression of a given target [13].

The present study aimed at differentiating the network activation effects induced by three DBS targets (STN, GPi, and NAc) by using computational approach, a multivariate pattern-classification analysis (MVPA) [23]. Total 21 datasets obtained from three groups of healthy swine (seven animals per each DBS target group) were used, in which underwent a block-designed DBS-fMRI experiment. In the analysis, MVPA was applied to classify BOLD activation pattern between groups with varying size of region-of-interests (ROI), in which the ROI was systematically expanded to cover whole brain from individual region. Our analysis was conducted on five functional networks, which have been believed to have critical connections to neurologic and neuropsychiatric disorders. MVPA can represent the BOLD response measured in a large population of voxels, as a single multidimensional data structure. By doing so, the pattern classifier can differentiate BOLD pattern of distributed nodes associated to a unique experimental condition, i.e., stimulation target [24, 25].

## METHODS AND MATERIALS

### Animal

Twenty-one healthy swine were implanted unilaterally (left hemisphere) with DBS electrodes in the subthalamic nucleus (STN), globus pallidus interna (GPi), or nucleus accumbens (NAc) (7 swine per target group, 35±5 kg). All study procedures were performed in accordance with the National Institutes of Health Guidelines for Animal Research (Guide for the Care and Use of Laboratory Animals) and approved by Mayo Clinic Institutional Animal Care and Use Committee.

### Animal surgery

Animals were initially sedated with the Telazol (5 mg/kg) and Xylazine (2 mg/kg) cocktail via intravenous bolus injection. The anesthesia was maintained with a medical grade gas mixture of O2 and N2o, and concentration of 1.5–2% isoflurane. Anatomical 3D MP-RAGE imaging was conducted in the General Electric (GE) Signa HDx 3.0 Tesla scanner. The imaging parameters are followed: repetition time (TR) = 11 ms, echo time (TE) = 5.16 ms, field of view (FOV) = 24 × 24 (mm), matrix size = 256 × 256, slice number = 128; 4-channel custom-built radio frequency (RF) coil. By using the COMPASS stereotactic planning software (COMPASS International Innovations, Rochester, MN), three dimensional coordinates (X, Y, and Z) for each target was identified and Leksell stereotaxic (Elekta, Stockholm, Sweden) coordinates were obtained from individual anatomical image volumes [26, 27]. A micro drive system (Alpha-Omega system) guided the implantation of The DBS electrode (Model 3389, Medtronic Inc.). The location of the DBS electrode was visually confirmed through MR-CT fusion images after a surgery.

### Stimulation design and fMRI acquisition

Stimulation of the STN, GPi, and NAc was conducted while gradient echo (GRE) echo-planar imaging (EPI) was conducting. The imaging parameters were as follows: TR = 3000 ms, TE = 34.1, flip angle (FA) = 90°, slice thickness = 2.4 mm, FOV = 15 × 15 mm, matrix size = 64 × 64. The total scanning time for each subject was 12 minutes 45 seconds. For each scan, the anesthetized was maintained with 1.2 ~ 1.4% isoflurane. After initial dummy scans for stabilizing the scanner, five stimulation-on and off periods were followed. Each block consisted of 6s of stimulation-on block and 120s of stimulation-off block. During DBS-fMRI experiment, all animal physiology including heart rate, respiration rate, end-tidal CO2, spO2, and rectal temperature was monitored and maintained in normal rage.

### Stimulation parameters

The stimulation was carried out with following parameters: bi-phasic mono-polar pulse train; voltage, 5 [V]; pulse frequency, 130 [Hz]; pulse width, 90 [μs] with the two of the four contacts on the lead, 0 (anode) and 1 (cathode). (Justification)

### fMRI data analysis

All analysis was performed by the AFNI software (Analysis of Functional NeuroImages) [28]. Individual EPI volumes were preprocessed as following orders: spike removal (3dDespike in AFNI), slice-timing correction (3dTshift), motion correction (3dvolreg) with six parameters of translation and rotation. Then individual volumes were co-registered to the Saikali pig brain atlas [29] with the cost function of the Local Pearson Correlation (LPC). Spatial smoothing (3 mm Gaussian Kernel Full-width-half-maximum) and temporal filtering (bandpass between 0.01 hz and 0.3 hz) was carried

After preprocessing, statistical mapping of stimulation-induced BOLD activation was followed by using general linear model. Baseline drift (six orders) and residuals of motion artifacts were included to regress out artefacts from time course. The beta coefficients of GLM with other statistics were estimated. To obtain the group-level activation maps with significant voxel clusters being determined for each of three groups, a statistical analysis (one-sample two-tailed t-tests; 3dttest in AFNI) was applied to the individual GLM estimation results. The statistical threshold level was set to *p* < .05 (t > 2.45; false discovery rate corrected [FDR]).

### Region-of-interests (ROI) generation

We first divided each subject’s brain into 73 regions according to the anatomical labels in the atlas [29] and then applied singular vector decomposition (SVD) on the voxels’ BOLD time courses for each of the 73 regions [30–32]. The GLM parameter estimation (beta coefficients) followed, yielding 73 beta coefficients in individual EPI datasets. Finally group-level statistical analyses were conducted (2^nd^ level analysis; within-group, one sample t-test for each of STN, GPi, and NAc group; between-group, two sample t-test for STN versus GPi, STN versus NAc, and GPi versus NAc).

For comparing BOLD activity in the large areas and networks among targets, the range of region-of-interests were expanded to a larger area (e.g., sensorimotor system) including anatomically and/or functionally homogenous regions (e.g., PMC and PreMC) according to the categorization of the large region [29] and networks (Table 1). Then between-group (two-sample) t-test was applied for comparing the group averaged BOLD responses as similar to the voxel-wise statistical analysis.

**Table 1.** Regions included in network-level analysis. Abbreviations can be found in Appendix A.

| Networks | Regions | Sub-regions |
| --- | --- | --- |
| Limbic/Reward Loop | Prefrontal Cortex | OFC |
|  | Cingulate Cortex | vaCC |
|  | Striatum | CD, NAc |
|  | Pallidum | GPI, SNc |
|  | Thalamus | vaTH, mdTH |
|  | Limbic | FX, HP, Am |
| Cognitive/Associative Loop | Prefrontal Cortex | DLPFC |
|  | Cingulate Cortex | daCC |
|  | Striatum | CD, NAc, Pu |
|  | Pallidum | GPI, SNc |
|  | Thalamus | vaTH, mdTH, vlTH |
| Basal-ganglia-thalamo-cortical (BGTC) Motor loop | Sensorimotor | PMC, PreMC |
|  | Striatum | Pu |
|  | Pallidum | GPI, SNc |
|  | Thalamus | vaTh; vlTH |
| Fronto-Cerebellar network | Prefrontal Cortex | DLPFC, aPFC, OFC |
|  | Cingulate Cortex | daCC, dpCC, vaCC, vpCC |
|  | Cerebellum | 12 Lobules |
| Prefrontal/Attentional network | Prefrontal Cortex | DLPFC, aPFC, OFC |
|  | Cingulate Cortex | daCC, vaCC |

### Analysis of network BOLD response: Multivariate pattern analysis (MVPA)

BOLD patterns of five functional networks known for their relevance to neurologic and psychiatric disorders were the focus of this study. Networks were identified in accordance with the literature: 1) the limbic/reward loop [33, 34]; 2) the cognitive/associative loop [35, 36]; 3) the basal-ganglia-thalamo-cortical (BGTC) motor loop [33]; 4) the fronto-cerebellar network [37], 5) the prefrontal/attentional network [38, 39]. To differentiate network BOLD pattern between the DBS target groups, we applied multivariate pattern analyses with a pattern classification algorithm, Fisher’s linear discriminant (LDA) [40]. We aggregated 73 regional beta coefficients into a multi-dimensional feature vector that represents the pattern of a network.

To perform the group classification and validate classification accuracy, we adopted a k-fold repeated cross-validation procedure with the random resampling method [41]. Seventy multidimensional vectors from comparing groups (7 subjects x 5 stimulation blocks per subject x 2 comparing groups) were randomly partitioned to an equal size of k blocks (k=10). Each of individual samples in k-1 blocks was labeled with a group label (STN, GPi, or NAc) for LDA training, and the remaining one block was tested by the LDA model without a group label. Training and testing procedures were repeated over 1000 times with random resampling. The accuracy of two-group classification (STN vs. GPi, STN vs. NAc, and GPi vs. NAc) for each network was calculated based on the number of samples that were correctly classified into their original group over the total number of samples.

For each of the five networks, the accuracy rates were averaged, and statistics (standard error of mean) were calculated. To test the significance of group-classification accuracy, one-sample t-tests were carried out for the accuracy rate of each network against the chance level. The empirical threshold of the chance level in a two group classification varies with the number of data being sampled [42]. Thus, chance was determined to be 59.8% in our study.

## RESULTS

### Group-level BOLD activation of STN, GPi, and NAc stimulation

Using the conventional voxel-wise analysis, we identified several cortical and subcortical clusters of voxels that registered significant BOLD changes relative to the pre-stimulation baseline. As shown in Fig. 1, numerous voxel clusters showed significant BOLD activation, reaching the threshold *(p* < .05; *t* > 2.45, FDR corrected).

**Fig. 1.**
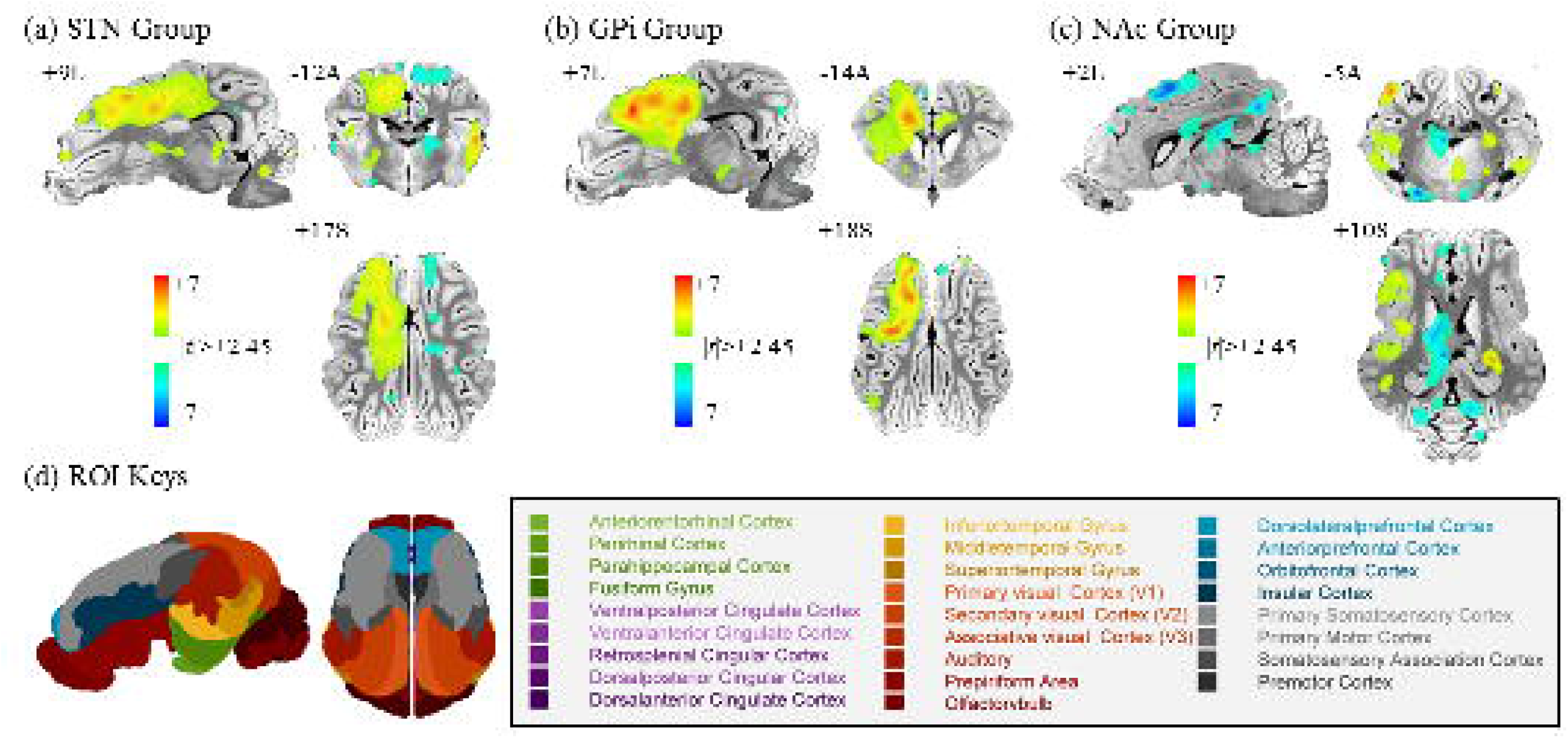
Group-level BOLD activation map of STN, GPi, and NAc DBS groups (n=7 for each group). The clusters of voxels showing a significant BOLD response to stimulation (5 [V], 130 [Hz], and 90 [μs]) are denoted in different colors. The statistical threshold of significance was set at *p* < .05 (one-sample two-tailed t-test, t[6] > 2.45). Stimulation of each of these basal ganglia structures elicited BOLD activation in multiple cortical and subcortical brain regions. Note that stimulation targets were located in the left hemisphere. (A) subthalamic nucleus stimulation group, (B) globus pallidus internus stimulation group, and (C) nucleus accumbens stimulation group. (D) Color-coded atlas of the domestic pig brain.

STN stimulation evoked a significant activation in the ipsilateral (left) primary motor cortex (PMC), primary somatosensory cortex (PSC), putamen (Pu), and in the contralateral (right) anterior prefrontal cortex (aPFC), insular cortex (IC), and prepiriform area (PPf). Similar to STN DBS, GPi DBS induced activation in the ipsilateral PMC, PSC, but also in the ipsilateral dorsal anterior and posterior cingulate cortex (dpCC; daCC), the dorsal lateral prefrontal cortex (DLPFC), and putamen. The BOLD patterns for STN and GPi are consistent with previously reported findings [16]. In contrast, NAc DBS reduced the BOLD response in PMC compared to the activation of motor areas noted in the STN and GPi groups. NAc DBS also induced a notable increase in the BOLD response in the striatum including the claustrum (Cl) including many other subcortical areas (reticular thalamic nucleus, and fornix). These results are in contrast with previous results, showing an increased BOLD response in the corresponding regions. However, the subtle spatial variation of the stimulation locus within the ventral striatum could result in a different BOLD pattern by way of increasing or decreasing BOLD. See the details of activated clusters in Table 1 and the list of abbreviations in Appendix A.

### Regional BOLD responses induced by DBS

Both STN and GPi stimulation induced a significant BOLD response *(p* < .001; *t*> 5.9; qFDR < .05) in the ipsilateral sensorimotor regions including PMC, PSC, and PreMC (Fig. 2). However, such modulatory effects between STN and GPi were different depending on the hemisphere, showing that STN reduced the BOLD response in PMC (Fig. 2a) in the contralateral (right) hemisphere, while GPi did not (Fig. 2b). NAc induced both an increase and decrease in BOLD activity in multiple regions, for example, increased responses in the IC, Cl and PPf (Fig. 2c), and decreased responses in the ipsilateral PMC and anterior ventral thalamus (avTH). Taken together with a voxel-wise analysis, the results of the regional BOLD analysis indicate that similar modulatory effects were induced between both STN and GPi, but these effects were distinct from NAc DBS. See the complete results for 73 regions in Supplementary Fig. S1.

**Fig. 2.**
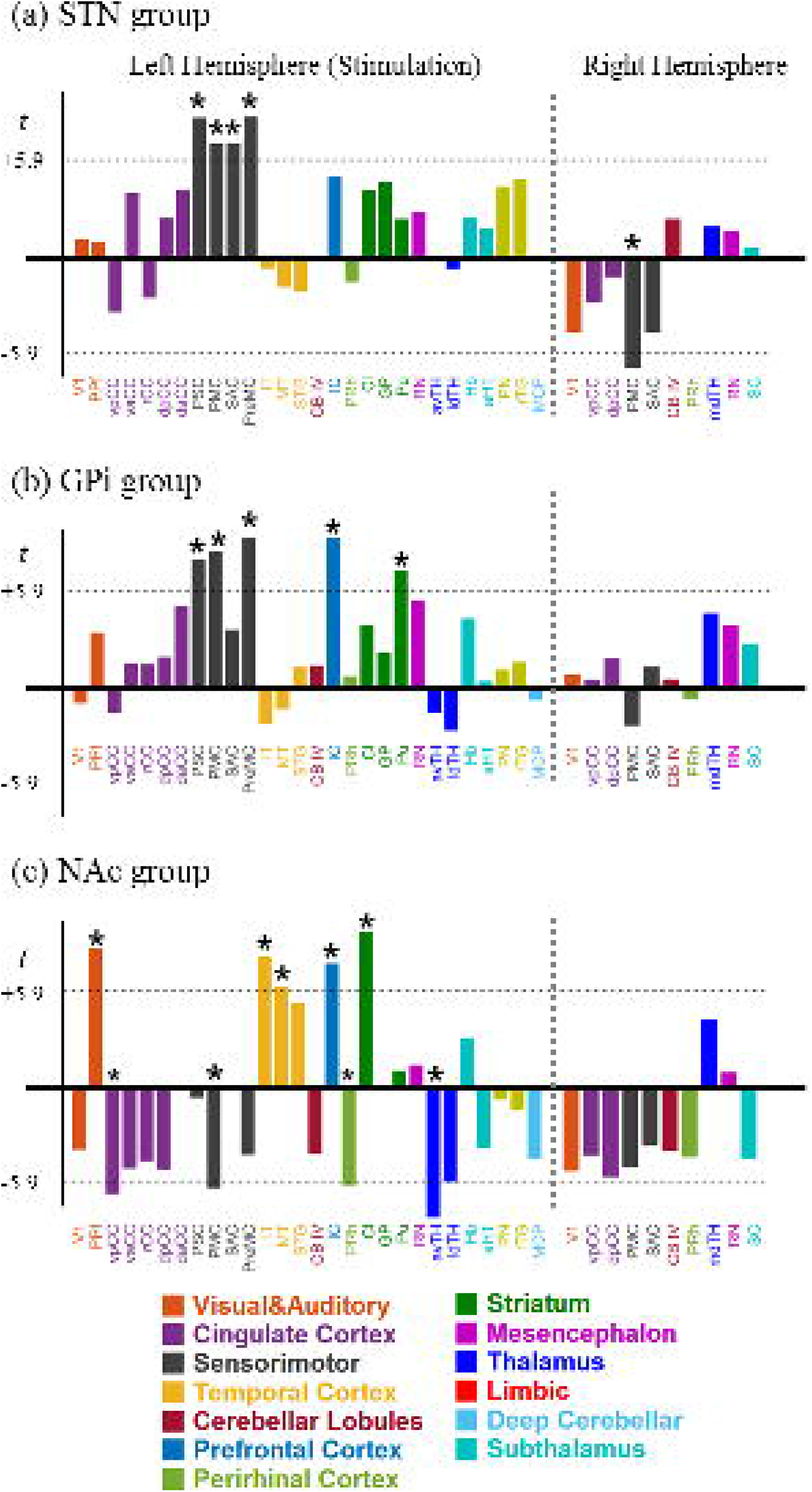
Group-level regional BOLD response of STN, GPi, and NAc DBS groups (n=7 per group). Both STN and GPi DBS evoked BOLD activation in motor regions, and NAc induced increase or decrease BOLD activity in multiple cortical and subcortical regions. Note that DBS applied on the left hemisphere. An individual subject’s whole brain was divided into 73 distinctive anatomical regions (30), and regional BOLD time courses were extracted. Parameter (beta coefficient) estimation was then conducted for each regional BOLD response. Group-level (n=7) analysis for each of 73 regions was followed. Bars indicate the t-value of corresponding regions, at least one of the three stimulation groups presented here. The asterisk (*) indicates BOLD response of the region reached statistical threshold at *p* < .001 (t[6] > 5.9, qFDR < .05). Abbreviations can be found in Appendix A. See Supplementary Fig. S1 for the complete results of 73 regions

We also conducted between-group statistical tests in order to examine distinct region-level BOLD responses (Supplementary Fig. S2). No significant regional differences were found between the STN and the GPi group, while the substantially different BOLD activity pattern was induced in the NAc group, compared to those of STN and GPi.

### Multivariate pattern analysis (MVPA): BOLD activity in large cortical areas

We expanded our region-of-interests (ROIs) to larger cortical areas, and conducted between-group pattern classification analyses through multivariate pattern analyses (MVPA). As a result, we found the presence of a distinguishable pattern among DBS targets in large cortical areas (Fig. 3). Specifically, STN and GPi stimulation induced a significant, distinct pattern in the ipsilateral visual, auditory system, striatum, and contralateral sensorimotor system (*p* < .05; chance level accuracy, 59.8%) (Fig. 3a). It is noteworthy that no distinctions were detected from the region level analysis (Supplementary Fig. S2). Between the STN and NAc groups, we noted that the patterns of several cortical and subcortical areas were different (Fig. 3b). Such a salient distinction was not unexpected, since from the regional BOLD analysis a large distinction was identified (Fig. 2c). GPi and NAc also induced distinctive BOLD patterns in large cortical areas, in which the classification performance was similar to that obtained from a comparison of STN and NAc. Taken together, the modulating effects between STN and GPi on large cortical and subcortical areas are analogous to each other, but different from that of NAc.

**Fig. 3.**
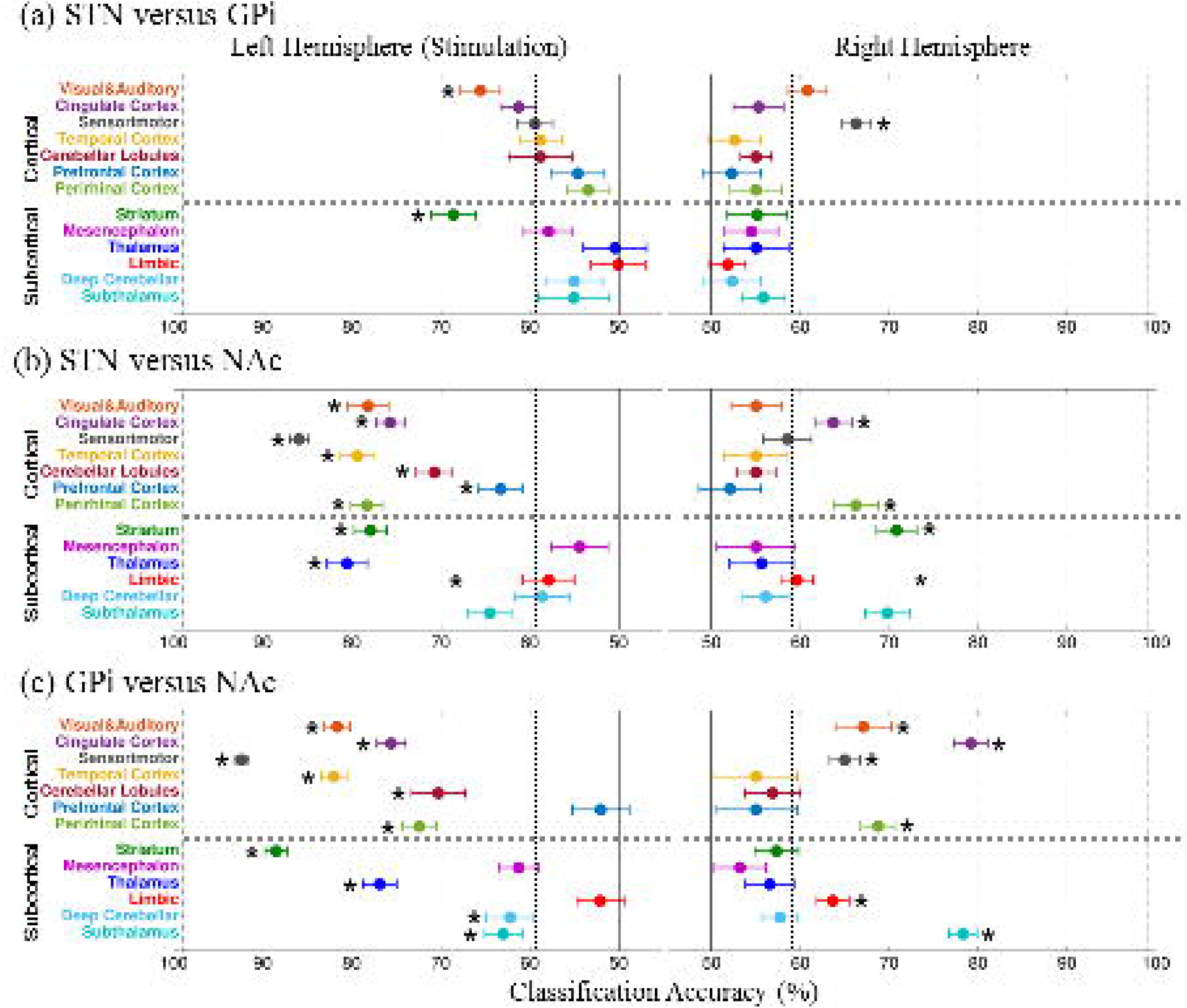
Between-group multivariate pattern classification results at large cortical and subcortical areas (p < .05; chance level, 59.3%). (A) STN versus GPi: Distinctive BOLD pattern was detected in visual & auditory, striatum, and contralateral sensorimotor cortices. (B) STN versus NAc: Most of the ipsilateral cortical and subcortical areas evoked distinctive patterns. (C) GPi versus NAc: Similar discrimination pattern was obtained with STN versus GPi classification. Dots indicate the accuracy of two-group classification with error bars of ±1 standard error of mean (SEM). The black vertical dashed line indicates the minimal correct classification rate (corresponding to chance level, 59.3%). Accuracy was calculated based on the number of individual samples that were correctly classified into corresponding group over the total number of samples. See details in Methods.

### Multivariate pattern analysis (MVPA): Network-level BOLD patterns

The pattern classification was applied to BOLD patterns of five disease-related brain networks. We found that the accuracy of classifying the BOLD pattern network between the STN and GPi group reached the level of statistical significance in the cases of the ipsilateral limbic/reward and the contralateral BGTC-motor networks (p < .05; chance level accuracy, 59.8%) (Fig. 4a). Between the STN and NAc group (Fig. 4b), all five ipsilateral networks (limbic, associative, motor, fronto-cerebellar, and prefrontal networks) evoked distinctive BOLD patterns. Similar classification results were obtained in the contralateral side as well. Notably, a higher classification accuracy in the ipsilateral BGTC-motor network (91 percent) and limbic network (88.7 percent) was obtained. GPi and NAc also induced distinctive network BOLD patterns. Similar with the results from large cortical areas (Fig. 3), this suggests the presence of analogous network-level modulatory effects between STN and GPi, but these were different from that of NAc.

**Fig. 4.**
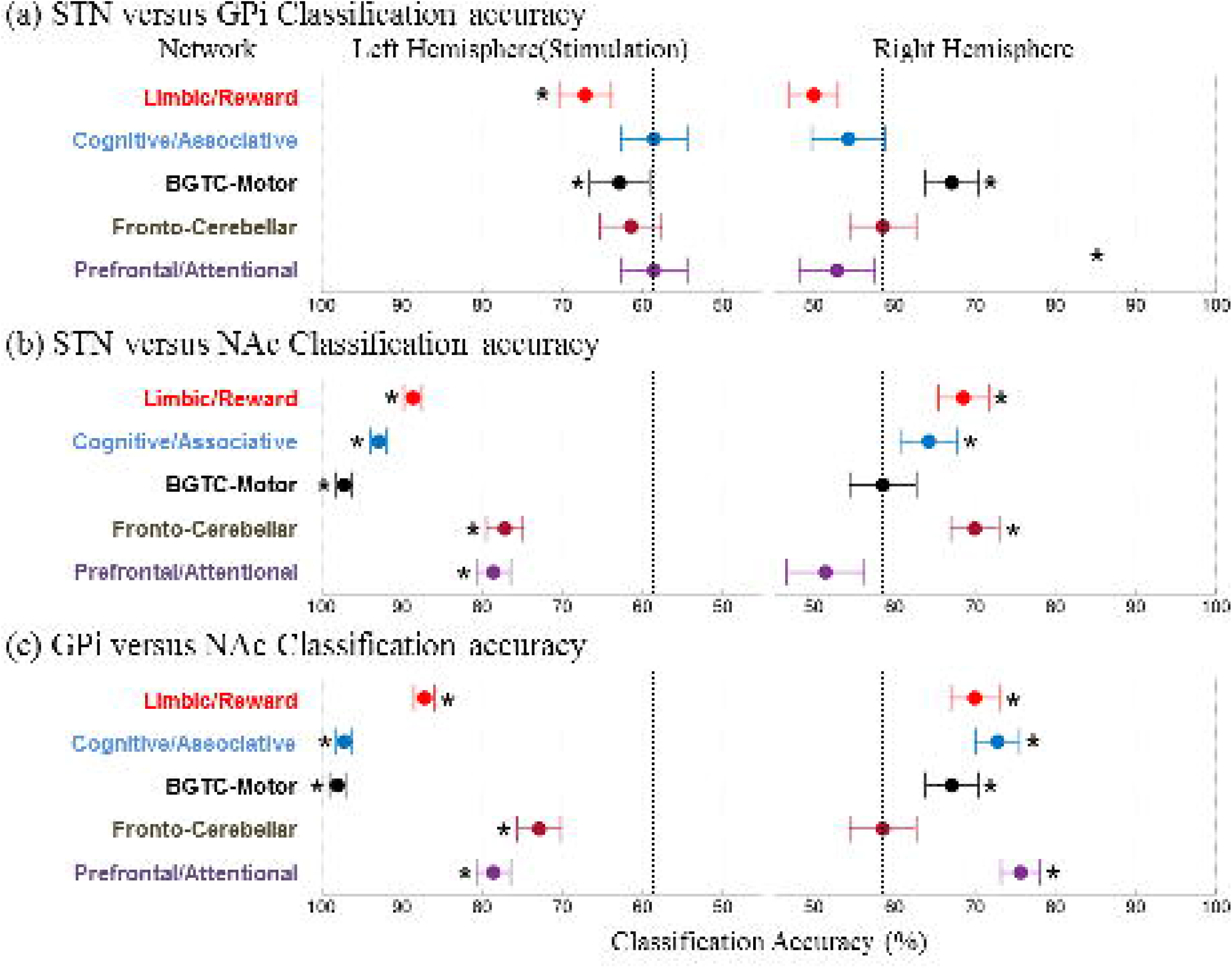
Between-group multivariate pattern classification results at five disease-related networks (*p* < .05; chance level, 59.3%). (A) STN versus GPi group: distinctive BOLD pattern was detected in the ipsilateral limbic and both ipsilateral and contralateral motor networks (B) STN versus NAc group: All five networks in ipsilateral and limbic, associative, and fronto-cerebellar networks evoked distinctive BOLD pattern (C) GPi versus NAc group: Similar results to (B) were obtained in the ipsilateral networks, but additional pattern distinction was found in the contralateral BGTC-motor network. Dots indicate the accuracy of two-group classification with error bars of ±1 standard error of mean (SEM). The black vertical dashed line indicates the minimal correct classification rate (corresponding to chance level, 59.3%). See the list of brain regions affiliated to each network in Table 1.

Although a univariate analysis detected some significant differences in contralateral BOLD patterns (Fig. 5a), MVPA was generally able to detect pattern distinctions in more networks (Fig. 5b). In particular, the large pattern distinction in the contralateral side of the STN and GPi group may be the result of a reduced STN DBS BOLD response in the contralateral hemisphere, whereas GPI did not (Fig. 2 and Supplementary Fig. S2).

**Fig. 5.**
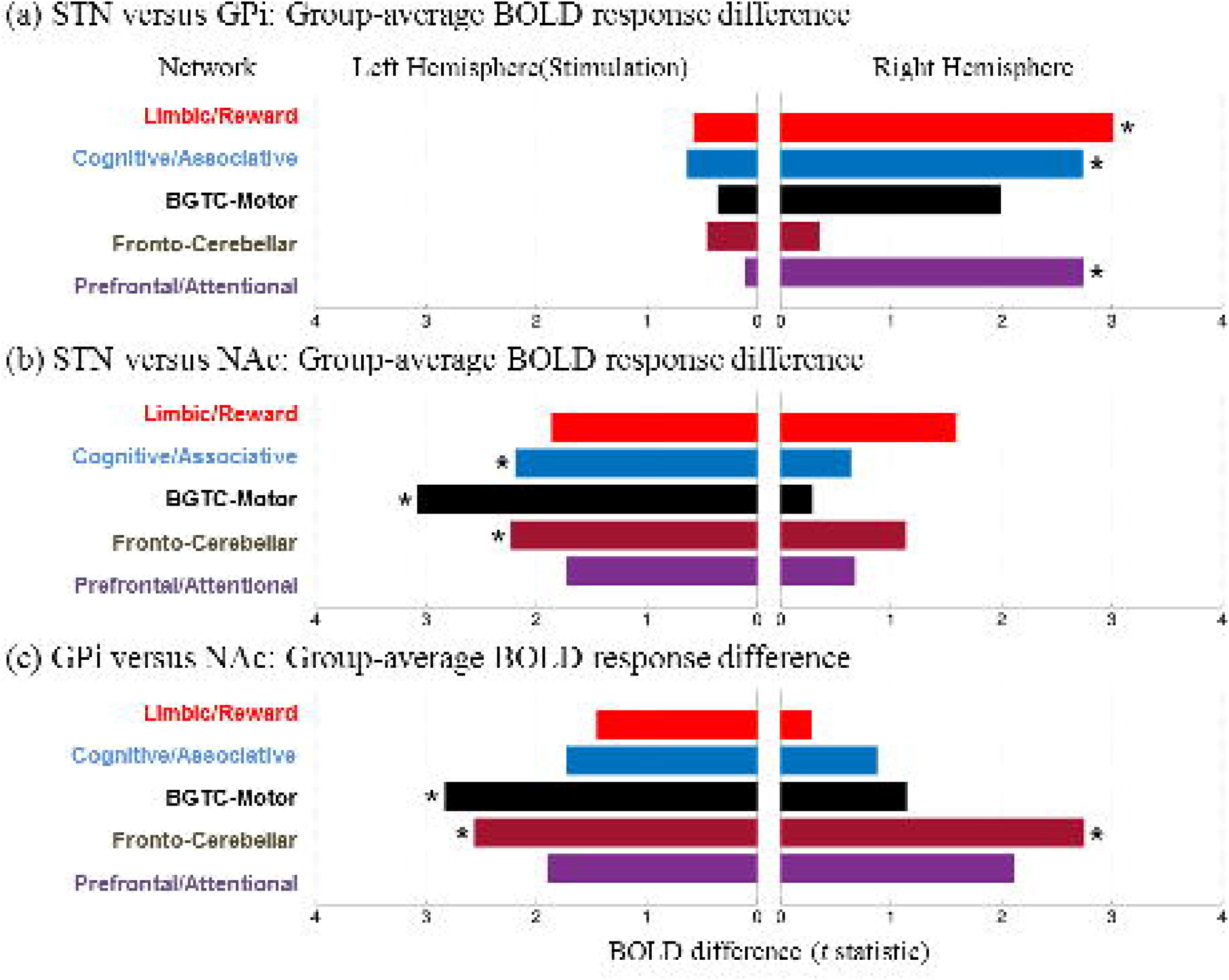
Between-group BOLD response comparison (two-sample *t*-test) at five disease-related networks (*t*[12] > 2.17; *p* < .05). Bars indicate the t-statistic of BOLD level difference between two comparing groups. (A) STN versus GPi group: Significance differences of BOLD level were found in the contralateral (right) hemisphere (limbic, associative, and prefrontal networks). (B) STN versus NAc: Ipsilateral three networks (associative, motor, and fronto-cerebellar) evoked the significantly different BOLD level. (C) GPi versus NAc: Both ipsilateral networks (motor and fronto-cerebellar) and contralateral (fronto-cerebellar) evoked different BOLD levels. See the list of brain regions affiliated to each network in Table 1.

Of note, our analysis demonstrates that MVPA was capable of differentiating the BOLD response between STN and GPi, suggesting that the multivariate approach provides a better sensitivity for discriminating between BOLD responses, particularly in larger ROI, whereas the univariate approach was less sensitive (Supplementary Fig. S3).

## DISCUSSION

The central findings of this study are that STN, GPi, and NAc DBS induced distinctive and characteristic BOLD responses on limbic, associative, and motor basal-ganglia-thalamo-cortical networks. Our results suggest the neurobiological and clinical significance of these findings in terms of 1) the target-specific clinical effectiveness of DBS may stem from its characteristic network modulation in large-scale brain circuits, and 2) a multivariate approach can provide a sensitive measurement for detecting subtle pattern differences registered in brain networks, which are not readily differentiated by conventional univariate voxel-wise analyses.

### Distinctive network effects of STN, GPi, and NAc on limbic, associative, and motor circuits

The multivariate pattern analysis (MVPA) revealed that stimulating three distinctive regions in the basal ganglia (BG) not only modulated disparate regions, but also induced distinctive modulatory patterns in brain networks. Both STN and GPI stimulation exert similar effects on motor-related regions [16], which has been clinically implicated as both targets for treating PD [43]. Both STN and GPi DBS are believed to eliminate or reduce pathological oscillations within BGTC-motor circuits, resulting in alleviating motor-related symptoms [14, 15, 33]. However, we noted that pattern distinctiveness was also observed along with brain networks (Fig. 4). Indeed some clinical reports indicated a different clinical efficacy of STN and GPi, for example, GPi-DBS attenuated chorea in Huntington’s disease [44], alleviated dystonia [45], but induced bradykinesia [43]. GPI DBS would be less effective for reducing levodopa medication [46] compared to that of STN DBS.

We also found that STN and GPi evoked distinctive patterns in the limbic network. STN is known to be a part of circuitry that is responsible for reward-related information processing, for example, an electrophysiological study in a non-human primate found that processing reward-related information may affect neural activity in the STN [47]. In the case of humans, a broad range of effects of STN DBS on reward-related behavioral and cognitive functions have been reported such as alleviating OCD symptoms [48–50]. It is possible that undesirable reward-related adverse effects in behavior and cognition are associated with STN-DBS [2, 43, 51], for example, pathological gambling [52], depression [53], and mania [54]. Additionally, it is conceivable that unilateral STN DBS has a bilateral effect, whereas the effect of GPi DBS may be limited to the ipsilateral side. Unilateral stimulation of STN was reported to be as effective as bilateral stimulation in alleviating symptoms of patients with PD [55], whereas it may not be the case for GPi [56].

Our data showed that STN and NAc DBS induced very distinctive modulatory effects across limbic, associative and motor networks. NAc DBS tended to reduce BOLD responses on regions, indicating its inhibitory effect on local neuronal activity. Early studies of NAc showed that stimulation suppressed motor execution such as eating behavior, and leg flexion [57–59], and vocalization [59, 60] in animal models [61]. However, other studies reported that NAc DBS in fact could increase the BOLD response on cortical areas [19, 20], presumably due to the subtle spatial displacement of a stimulation locus [19] within complex striatal substructures [62–64].

### Clinical applications of multivariate approach to DBS-fMRI data

We highlight here that multivariate pattern analysis (MVPA) has better discriminative characteristics for comparing overlapped and sparsely distributed BOLD patterns, particularly in the network level of a brain. Many recent studies have taken advantage of pattern discrimination sensitivity in a multivariate approach, showing that discrete BOLD patterns were associated with differing visual tasks [65, 66], different emotions [67], and memory retrieval processes [68]. In addition, it was able to diagnose a psychiatric disorder (e.g., depression) [69, 70], to identify the distinctive brain patterns of heroin-dependent individuals from healthy subjects [71], to predict symptom severity in autism [72], and to delineate subgroups of schizophrenia patients [73]. We demonstrated the existence of target-dependent discrete network patterns within complex thalamocortical circuits, which highly likely related to therapeutic effects for movement and neuropsychiatric disorders [33]. This will improve our understanding of the DBS impact on brain circuits, which eventually helps to optimize the current DBS therapy.

### The potential factors that impact BOLD patterns

It should be noted that anesthesia and the electrophysiological properties of stimulation parameters in our study may result in a biased interpretation of BOLD activation. The level of anesthesia [74] and the type of anesthetic agent used [75, 76] can have different attenuation effects on BOLD signal changes [77]. Because the anesthetic used herein, in general, reduced the BOLD response, it is likely that the brain responses observed in our study were underestimated compared to the non-anesthetized condition. However, we believe that our experimental design has already been validated by previous studies in terms of inducing a robust and stable BOLD response [16, 20, 78], thus the impact of anesthesia on the global brain less likely to mislead the current results of the network-level analysis. Alternatively it is also conceivable that the BOLD activation might have been overestimated when compared to a study which used a long-term steady state stimulation [79–81]. The short-term bursts of pulse train (6 sec) and our stimulation parameters generally could evoke a larger, acute BOLD response [82]. Finally, the potential interaction between stimulation duration and anesthetic agent remains uncertain, thus future studies are needed to elucidate such complications.

Since our subjects were healthy, BOLD response patterns in healthy brains may not be replicated in a diseased-state brain, since such a disease may alter network connectivity. For future studies, it is therefore necessary to test the classification performance of MVPA in disease-state brains to differentiate target-dependent DBS BOLD response (e.g., MPTP animal model of Parkinsonism).

Although our study did not show a correlation between direct behavior and stimulation-induced network effects, in part, due to the limitations of an animal study, yet we believe that our approach can be used to explain or predict the behavioral manifestation, providing correlates to DBS-induced network effect.

DBS electrode creates metal-induced artifacts in the MR image, thus the BOLD effects near the electrode lead could be underestimated (Supplementary Fig. S5). Although we excluded voxels near the electrode lead in our analysis, the extent and magnitude of artifact still remains uncertain. Future studies should quantitatively measure the potential impact on lead artifacts [83].

Our data showed that these DBS targets have the similar and dissimilar BOLD pattern features across brain networks, which were not readily detected in the previous studies that relied on the use of a conventional univariate approach. These results improve our understanding of neural network effects of differing targets on the therapeutic and adverse outcomes of DBS. Furthermore, our proposed approach represents a promising tool for identifying and distinguishing previously unidentified network-level brain activity, which correlates therapeutic efficacy in clinical disciplines. The target-specific clinical effectiveness of three DBS targets likely stems from its characteristic network modulation in large-scale brain circuits.

## ACKNOWLEDGEMENTS

This work was supported by The Grainger Foundation and by the National Institutes of Health (R01 NS 70872 awarded to K.H.L.). The authors thank Dr. Penelope S. Duffy for helpful discussion.

## APPENDIX A: Abbreviations

aPFC: anterior prefrontal cortex
Am: Amygdala;
Au: primary auditory cortex;
CB III: cerebellar lobule III;
CB IV: cerebellar lobule IV;
daCC: dorsal anterior cingulate cortex;
dpCC: dorsal posterior cingulate cortex;
rCC: retrosplenial cingulate cortex;
vaCC: ventral anterior cingulate cortex;
Cd: caudate;
CD: caudate nucleus;
Cl: claustrum;
DLPFC: dorsal lateral prefrontal cortex;
FX: fornix;
GP: globus pallidus;
GPi: globus pallidus interna;
Hb: habenular nuclei;
HP: hippocampus;
aHT: anterior hypothalamic area;
IT: interior temporal gyrus;
IC: insular cortex;
MCP: medial cerebellar peduncle;
MT: middle temporal gyrus;
NAc: accumbens nucleus;
OFC: orbitofrontal cortex;
PreMC: premotor cortex;
PMC: primary motor cortex;
PN: pontine nucles;
PPf: prepiriform area;
PRh: perirhinal cotex;
PSC: primary somatosensory cortex;
Pu: putamen;
RN: red nucleus;
SAC: sensory association cortex;
SC: superior colliculus;
SNc: substantia nigra pars compacta;
STG: superior temporal gyrus;
rTG: renticulotegmental nucleus;
avTH: anterior ventral thalamic nucleus;
ldTH: lateral dorsal thalamic nucleus;
mdTH: mediodorsal thalamic nucleus;
rTH: reticular thalamic nucleus;
vaTh: ventral anterior thalamic nucleus;
vpCC: ventral posterior cingulate cortex
vlTh: ventral lateral thalamic nucleus;
V1: primary visual cortex;

---

**Supplementary Fig. 1. Group-level regional BOLD response of STN, GPi, and NAc DBS groups in 73 brain regions (n=7 for each group).**

**Supplementary Fig. 2. Between-group BOLD response comparisons at the regions**. DBS-evoked BOLD responses between stimulation groups were compared between corresponding anatomical regions shown in (A) STN versus GPi group, (B) STN versus NAc group, and (C) GPi versus NAc group (n=14, two-sample, two tailed *t* test). Bars indicate the t-value of corresponding regions. The asterisk (*) indicates BOLD response of the region reached the statistical threshold at *p* < .001 (t > 3.9, qFDR < .05). Abbreviations can be found in Appendix A.

**Supplementary Fig. 3. Between-group BOLD response comparisons at the large cortical and subcortical areas**. DBS-evoked BOLD responses between stimulation groups were compared between corresponding areas shown in (A) STN versus GPi group, (B) STN versus NAc group, and (C) GPi versus NAc group (n=14, two-sample, two tailed *t* test). Bars indicate the t-value of corresponding cortical or subcortical area. The vertical dashed line denotes the statistical threshold of *p* < .001 (t > 3.9, qFDR < .05). Results for cortical and subcortical regions are shown above and below the horizontal dashed line, respectively. The brain regions and their affiliation to large cortical and subcortical areas can be found in [29].

**Supplementary Fig. 4. Between-group multivariate pattern classification results mapped onto the pig brain model**. Differing intensity of color indicates the group classification accuracy. The brain regions and their affiliation to large cortical and subcortical areas can be found in [29].

**Supplementary Fig. 5. Susceptibility artifact in echo-planar images (n=7) of DBS electrode lead.** The width/length of signal drop out area can be seen in (A) STN, (B) GPi, and (C) NAc DBS group. EPI images are shown in which the geometric image slice distortion is negligible along with readout (RO) dimension, while influential through the phase-encoding (PE) dimension. To minimize the distortion, the PE dimension was chosen in the direction where the length of the pig brain was the shortest.

