## Supplementary figures and images for "Multivariate pattern classification on BOLD activation pattern induced by deep brain stimulation in motor, associative, and limbic brain networks"

### Supplementary Fig. S1

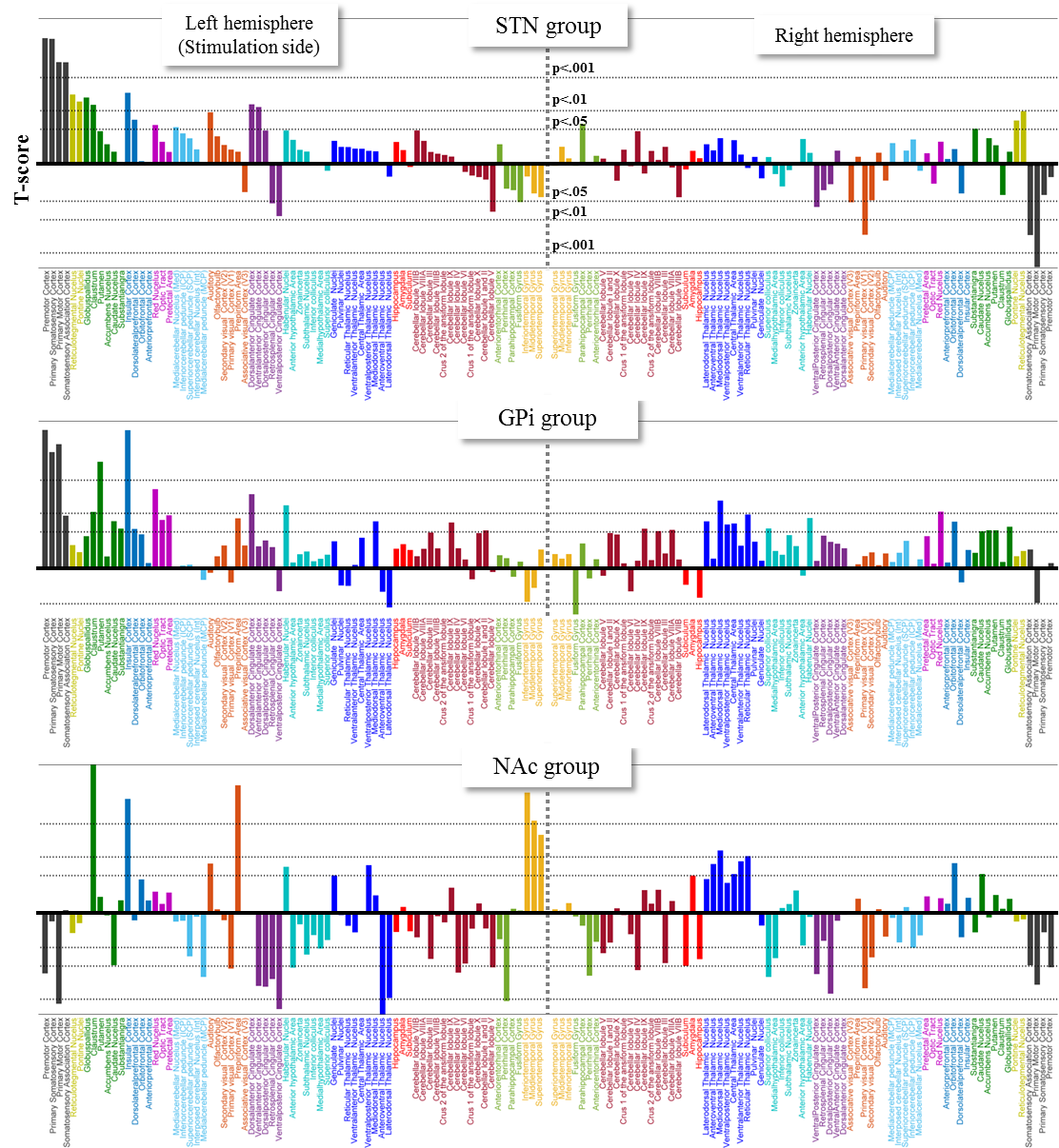

### Supplementary Fig. S2

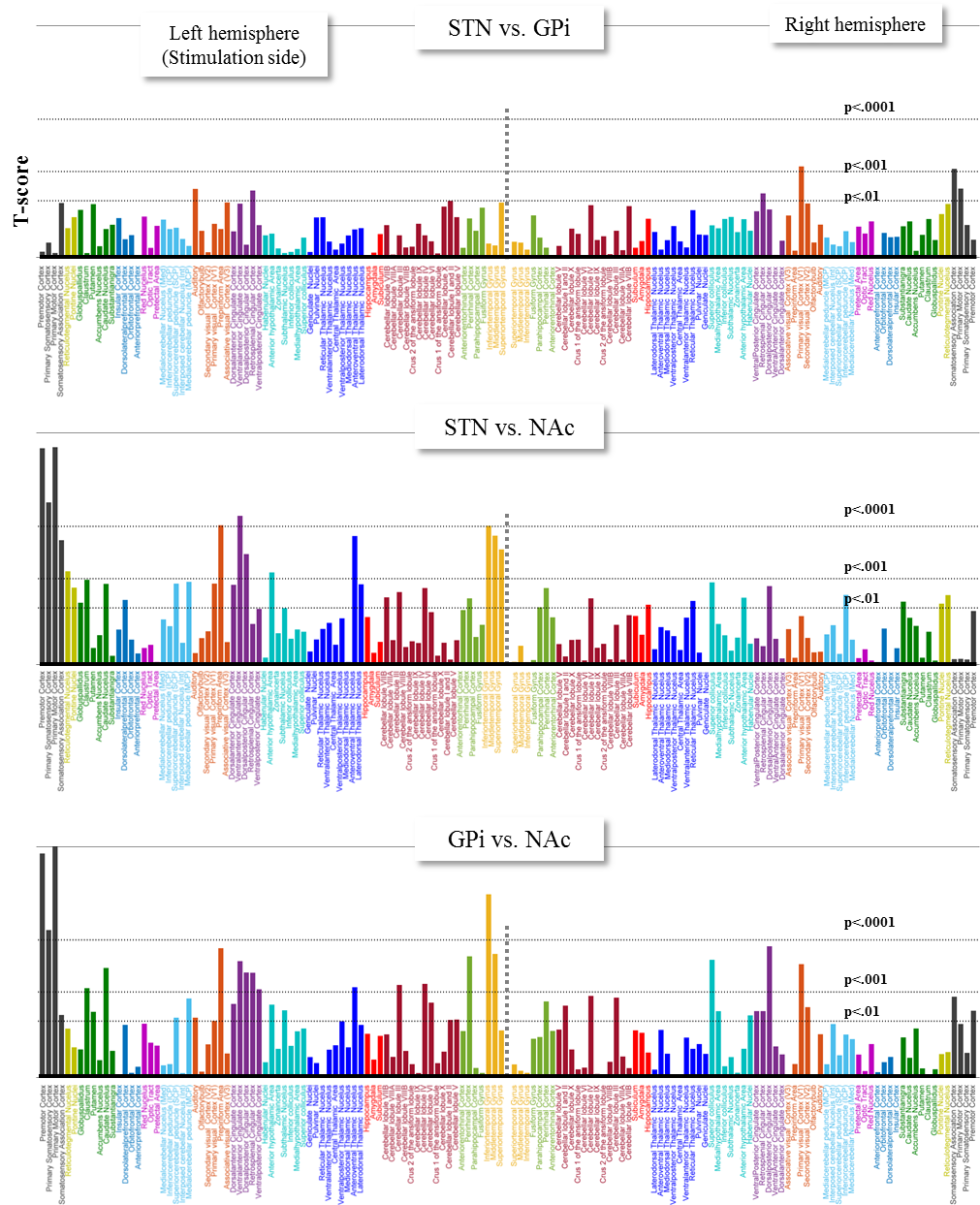

### Supplementary Fig. S3

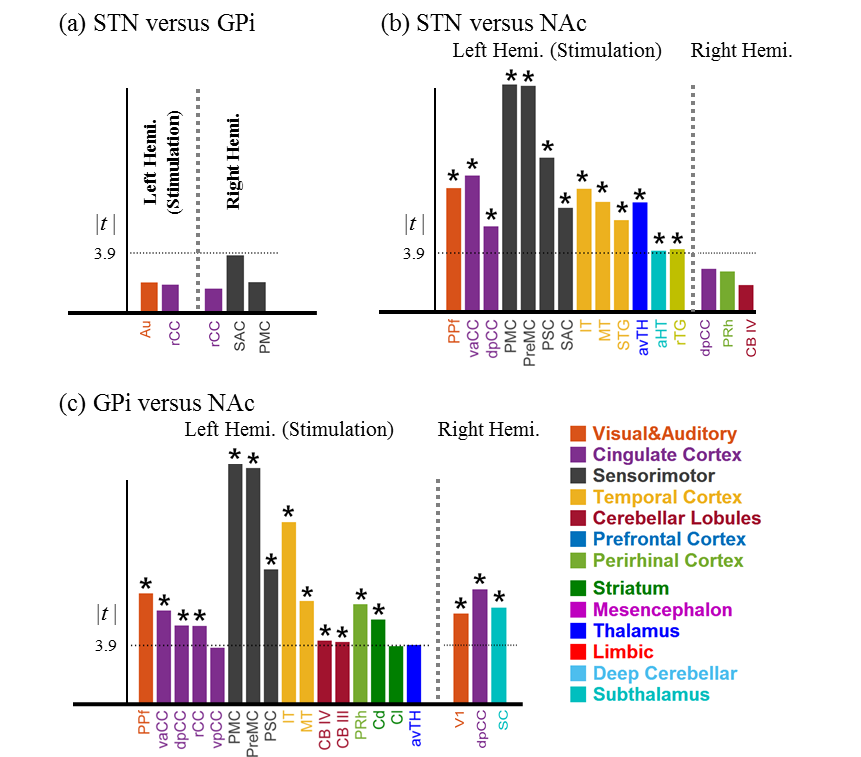

### Supplementary Fig. S4

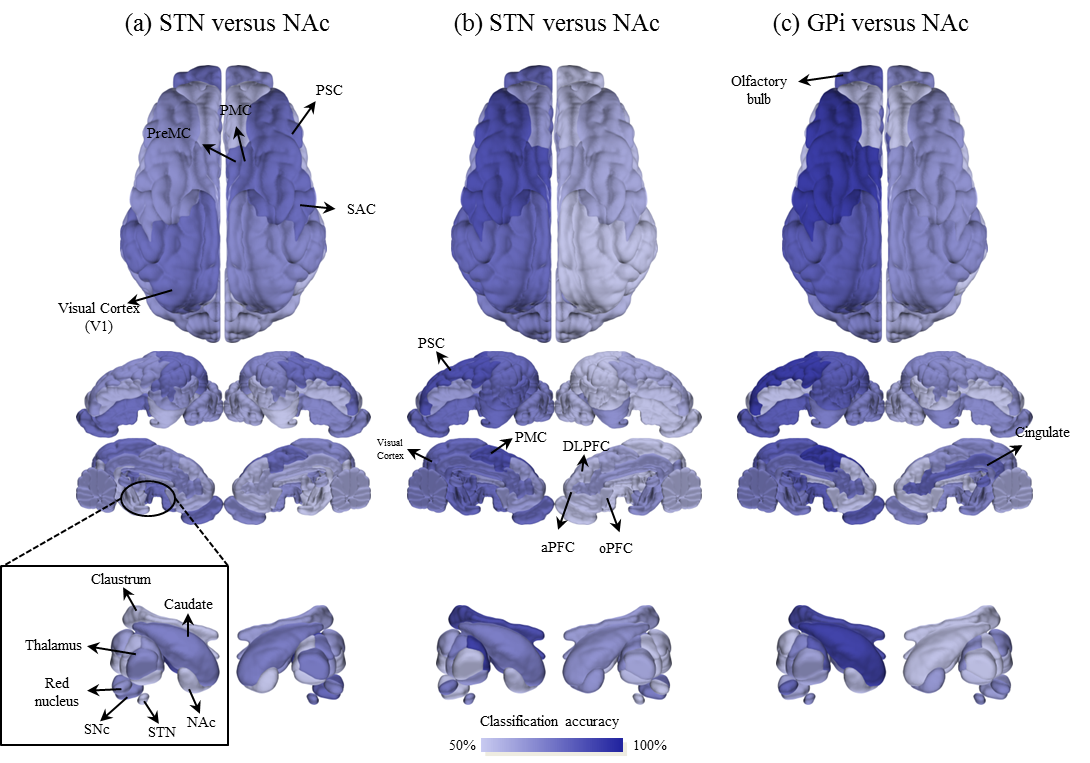

### Supplementary Fig. S5

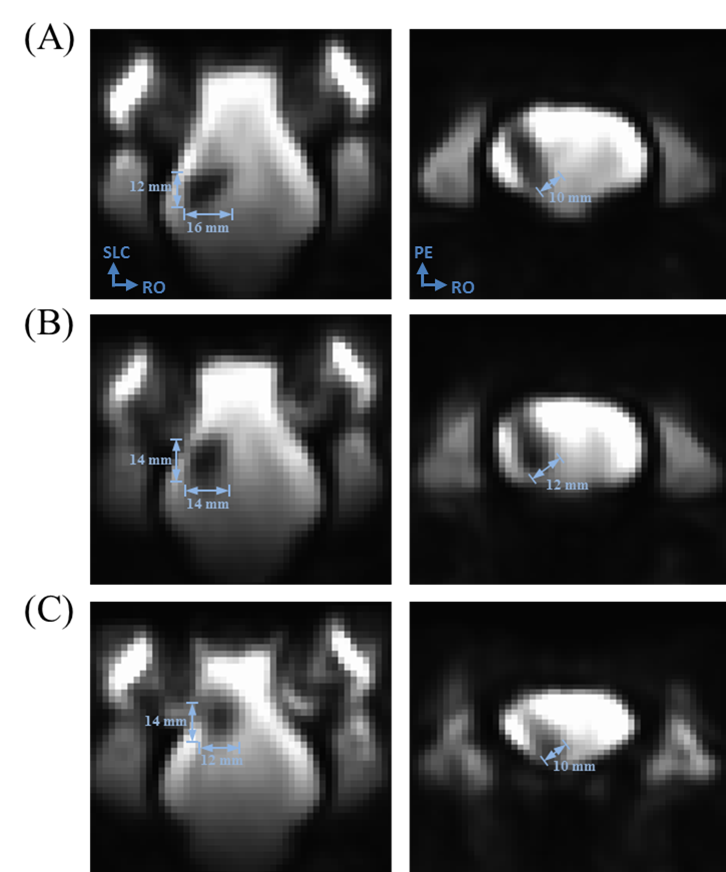
